# L-Phenylalanine restriction amplifies boron neutron capture therapy efficacy through increased L-boronophenylalanine uptake and induces activating transcription factor 4 stress response in tumor cell lines

**DOI:** 10.1101/2025.11.27.690635

**Authors:** Yuki Tamari, Rie Saba, Takushi Takata, Hiroyuki Kimura, Mao Takita, Kenta Yashiro, Minoru Suzuki, Hideya Yamazaki, Kei Yamada

**Affiliations:** Department of Radiology, Kyoto Prefectural University of Medicine, Kyoto, Japan; Institute for Integrated Radiation and Nuclear Science, Kyoto University, Osaka, Japan; Department of Medical Information Engineering, Research Promotion Unit, School of Medical Science, Fujita Health University, Aichi, Japan; Division of Anatomy and Developmental Biology, Department of Anatomy, Kyoto Prefectural University of Medicine, Kyoto, Japan; Division of Probe Chemistry for Disease Analysis, Research Center for Experimental Modeling of Human Disease, Kanazawa University, Ishikawa, Japan; Shimadzu Techno-Research Inc., Kyoto, Japan

**Author notes:** To whom correspondence may be addressed: **Yuki Tamari**, PhD, Department of Medical Information Engineering, Research Promotion Unit, School of Medical Science, Fujita Health University, 1-98 Dengakugakubo, Kutsukake-cho, Toyoake, Aichi 470-1192, Japan. **Rie Saba**, PhD, Department of Radiology, Kyoto Prefectural University of Medicine, 465 Kajii-cho, Kawaramachi-Hirokouji, Kamigyo-ku, Kyoto 602-8566, Japan. These authors contributed equally (Yuki Tamari, Rie Saba). **Authors responsible for statistical analysis:** Yuki Tamari,. Rie Saba,.

**Keywords:** Boron neutron capture therapy (BNCT), L-boronophenylalanine (BPA), L-phenylalanine, amino acid restriction, tumor cell line

## Abstract

Boron neutron capture therapy (BNCT) relies on the selective uptake of boron-10 compounds by tumor cells. L-boronophenylalanine (BPA) serves as a key carrier, and enhancing its accumulation is critical for improving BNCT efficacy. In this study, we investigated the effects of L-phenylalanine (Phe) restriction on the tumor cell lines SAS, U87-MG, PANC-1, and A375, and the immortalized keratinocyte line HaCaT on modulating BPA uptake and associated cellular responses. Quantitative analysis using inductively coupled plasma atomic emission spectroscopy (ICP-AES) showed that 24 h Phe restriction increased BPA uptake in the SAS, U87-MG, and PANC-1 cell lines. Colony formation assays confirmed enhanced sensitivity to neutron irradiation in these cells. RNA sequencing indicated that Phe restriction activated the integrated stress response downstream of activating transcription factor 4 (ATF4), although this pathway was not directly linked to increased BPA uptake. The L-type amino acid transporter 1 (LAT1)/4F2 heavy chain (4F2HC) complex was identified as the exclusive transport route for BPA. Liquid chromatography-tandem mass spectrometry (LC-MS/MS) analysis showed that Phe restriction altered intracellular levels of amino acids that serve as LAT1/4F2HC exchange substrates, suggesting a metabolic basis for enhanced BPA transport. Our results reveal that Phe restriction enhances BPA uptake and BNCT efficacy in a cell line-dependent manner, likely through the modulation of amino-acid metabolism. Therefore, targeted amino-acid manipulation prior to BNCT may represent a promising strategy to improve therapeutic outcomes.

## Introduction

Boron neutron capture therapy (BNCT) is a type of radiotherapy that exploits the ability of boron-10 (^10^B) to absorb thermal neutrons with high efficiency, resulting in nuclear decay to helium and lithium nuclei. The average range of these two heavy ion particles is 7 µm, which is shorter than the diameter of a single cell. Leveraging this property, BNCT can selectively destroy tumor cells that accumulate ^10^B-containing boron drugs during thermal or epithermal neutron irradiation. Clinically, BNCT has been performed at research reactors, and cyclotron-based neutron sources have been developed to facilitate its wider application (1–3).

L-boronophenylalanine (BPA), a boronated L-phenylalanine (Phe) analog, is the only BNCT agent approved under the Japanese national insurance system. The active uptake of amino acids via the high-level expression of the L-type amino-acid transporter 1 (LAT1)/4F2 heavy chain (4F2HC) complex, a major pathway for BPA incorporation, significantly contributes to tumor cell proliferation (4). The expression of this complex in normal tissues is limited (5, 6), leading to the selective uptake of BPA via LAT1 in tumor cells. However, expression of other amino-acid transporters has been reported to reduce the tumor-to-normal (T/N) ratio of BPA accumulation (7). Therefore, strategies to further increase tumor-selective BPA uptake are needed to optimize BNCT delivery.

Amino-acid restriction has been shown to inhibit tumor cell proliferation *in vitro* and improve survival in tumor-bearing animal models (8–10). Phe is an essential amino acid for protein synthesis and tyrosine biosynthesis, and a low-Phe diet is commonly used to treat patients with phenylketonuria (PKU) (11). Previous research has demonstrated that pretreatment with Phe overload increases the T/N ratio of BPA (12). However, the effect of Phe restriction on BNCT efficiency has not been established. Understanding how Phe restriction modulates BPA uptake by tumor cells is crucial for optimizing BNCT.

In this study, we aimed to evaluate the effects of Phe restriction on BPA accumulation and neutron irradiation efficacy and to investigate the associated cellular responses and underlying molecular mechanisms.

## Materials and methods

### Cell culture

SAS (13), U87-MG (14), PANC-1 (15), A375 (16), and HaCaT (17) cells were cultured in standard media (DMEM/F12 [08460-95; Nacalai Tesque, Kyoto, Japan], EMEM [056-08385; FUJIFILM Wako Pure Chemical Corporation, Osaka, Japan], and DMEM [08458-45; Nacalai Tesque]) supplemented with 10% fetal bovine serum (FBS; SH30396.03; Hyclone, Logan, UT, USA) and 1% penicillin-streptomycin solution (26253-84; Nacalai Tesque). The Phe-free medium (Gmep Inc., Kurume, Fukuoka, Japan) contained 10% dialyzed FBS (S-FBS-NL-065; Serana, Brandenburg, Germany) and 1% penicillin-streptomycin. Phe (13039-62, Nacalai Tesque) was added (2 µM–10 mM) as necessary.

### BPA uptake

^10^B-enriched BPA powder was obtained from Interpharma Praha (Prague, Czech Republic). A 3% (w/v) BPA/fructose solution was prepared by adding 0.30 g of BPA powder to a mixture of 1.52 mL of 1N NaOH and 8.0 mL of distilled water. Subsequently, 0.31 g of fructose was added, and the mixture was stirred until dissolved. Hydrochloric acid (HCl; 1 mol/L) was added to adjust the pH to 7.6. The 3% BPA-fructose solution was added to the cell culture medium at a final concentration of 10–200 µg/mL.

### Inductively coupled plasma–atomic emission spectrometry

The BPA-treated cell lines were washed with phosphate-buffered saline (PBS) and incubated overnight in the dark after the addition of 0.5 mL of 1N nitric acid. Distilled water (3.5 mL) was added to each sample and centrifuged at 2000 rpm for 10 min. The samples were filtered using a syringe filter (25 mm, 0.22 µm; Hawach Scientific, Xi’an, China) and analyzed using inductively coupled plasma–atomic emission spectrometry (ICP-AES; ICPE-9000, Shimadzu, Kyoto, Japan). The intracellular boron concentration was calculated using the values obtained from the ICP-AES and the following equation:

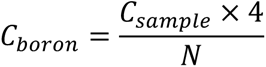

where C_boron_ is the boron concentration per 1 × 10^6^ cells (µg/1 × 10^6^), C_sample_ is the value obtained from the ICP-AES (µg/mL), and N is the total number of cells in the sample.

### Colony formation assay following neutron irradiation

The samples were irradiated in a research reactor at the Kyoto University Institute for Integrated Radiation and Nuclear Science. The cell lines treated with 3% BPA-fructose solution were trypsinized after 90 min and suspended in Phe-free medium containing 3% BPA-fructose solution. The cell suspension (0.5 mL) was poured into 1.5-mL tubes, attached to Li-6-enriched LiF tiles (six on the top and six on the bottom), and irradiated with neutrons. Two gold foil samples were attached to the top and bottom (front and back) of the tube, and two thermoluminescent dosimeters (TLDs) and an additional sheet of gold foil were placed at one location. The thermal neutron fluence and gamma-ray dose were measured and calculated using the gold foil and TLDs. The irradiated cells were immediately seeded (non-irradiated cells: 100; irradiated cells: 1,000–50,000) into 60-mm dishes and cultured in a control medium under 5% CO_2_ at 37°C. Cells were washed with PBS and fixed with ethanol after 14 days, stained with Giemsa stain, and colonies comprising at least 50 cells were counted after drying. The surviving fraction was calculated using the plating efficiency.

### RNA sequencing

#### Library preparation and sequencing

Total RNA was extracted from the cells using the RNeasy Mini Kit (74104, Qiagen, Venlo, Netherlands) and treated with recombinant DNase I (2270A, Takara, Kyoto, Japan). The RNA concentration was determined using a NanoDrop Lite spectrophotometer (A30955JP, Thermo Fisher Scientific, Madison, WI, USA). RNA sequencing (RNA-seq) libraries were prepared using the TruSeq Stranded mRNA kit (Illumina Inc., San Diego, CA, USA) following the manufacturer’s protocol. The libraries were sequenced on the NovaSeq 6000 and NovaSeq X Plus platforms (Illumina Inc.) using 101 bp and 151 bp single-end reads, respectively. Library preparation and sequencing were performed at the NGS Core Facility of the Research Institute for Microbial Diseases, Osaka University.

#### Data analysis

Raw sequencing reads were processed using Trimmomatic (18) to remove adapter sequences and assessed using FastQC (19). The processed reads were aligned to the human reference genome (hg38; https://www.ncbi.nlm.nih.gov/datasets/genome/GCF_000001405.26/) using STAR (20). Read counts per gene and transcripts per million (TPM) values were determined using RSEM (21). Differentially expressed genes (DEGs) with p-values < 0.05 and fold changes ≥ 2 between the Phe (+) and (–) conditions were identified using R (version 4.1.3; https://cran.r-project.org). Gene Ontology (GO) analysis of the DEGs was performed using Metascape (https://metascape.org/gp/index.html#/main/step1) (22). The RNA-seq data obtained in this study were deposited in the Gene Expression Omnibus database (http://www.ncbi.nlm.nih.gov/geo) under the accession numbers GSE279290, GSE279291, GSE279292, and GSE279293.

#### Immunofluorescence

Cells (5 × 10^4^ cells/well) were plated in 24-well plates and cultivated in normal medium for 24 h before incubation in Phe-deprived medium for 6 h. Subsequently, cells were fixed in ice-cold 4% paraformaldehyde (PFA)/PBS for 15 min before washing with PBS and treating with 3% bovine serum albumin (01860-36, Nacalai Tesque)/PBS containing 0.1% Triton X-100 (35501-15, Nacalai Tesque) (PBST) for 30 min. The primary and secondary antibody reactions were performed at 4℃ overnight and 25°C for 1 h, respectively. The following antibodies were used and diluted in PBST: rabbit anti-ATF4 (1:400 dilution; #11815, Cell Signaling Technology, Danvers, MA, USA) and Alexa Fluor 488-conjugated F(ab’)2 fragment donkey anti-rabbit IgG (1:2000; 711-546-452, Jackson ImmunoResearch, Philadelphia, PA, USA). DNA was stained with 4′,6-diamidino-2-phenylindole (DAPI; 11034-56, Nacalai Tesque). Fluorescent images were obtained using a BZ-X710 microscope (Keyence, Osaka, Japan).

### Amino-acid quantification by liquid chromatography-tandem mass spectrometry

#### Sample preparation

Cells were dissociated using 0.25% trypsin-EDTA, washed three times with ice-cold PBS, collected in 1.5-mL tubes, frozen in liquid N_2_, and stored at –80℃ until extraction for liquid chromatography-tandem mass spectrometry (LC-MS/MS) analysis at Shimadzu Techno-Research, Inc. (Kyoto, Japan). Each sample contained 1.0–2.0 × 10^6^ cells. Cell pellets were treated with 200 µL of methanol (Kanto Chemical Co., Inc., Tokyo, Japan) and 100 µL of H_2_O. Subsequently, 1000 µM of 2-morpholinoethanesulfonic acid (Tokyo Chemical Industry Co., Ltd, Tokyo, Japan) was added, and samples were homogenized. Homogenates were centrifuged at 10,000 rpm for 5 min at 4℃. Supernatants were dehydrated, reconstituted in 100 µL of H_2_O, filtered, and diluted for LC.

#### LC-MS/MS analysis

LC was performed using a Discovery HS F5-3 HPLC column (587503-U; Merck, Rahway, NJ, USA) and a LC-30A system (Shimadzu Co.). MS/MS was performed using the LC-MS/MS Method Package for Primary Metabolites Ver. 3 (Shimadzu Co.; https://www.shimadzu.com/an/products/liquid-chromatograph-mass-spectrometry/lc-ms-software/lcmsms-method-package-for-primary-metabolites/index.html) with a LCMS-8060NX (Shimadzu Co.).

#### Peak analysis

The internal correction value for each compound was calculated by dividing its peak area by that of the internal standard (IS) using the following formula:

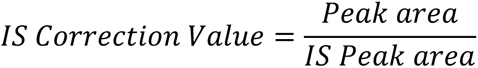

The peak area ratio was calculated by dividing the IS correction value of the Phe (+) sample by that of the Phe (–) sample as follows:

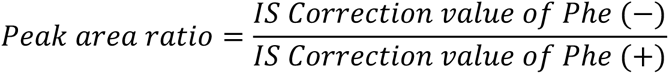

## Results

### Phe deprivation increases BPA uptake in tumor cell lines

To evaluate the effect of Phe restriction on BPA uptake, we measured the cellular ^10^B concentrations using ICP-AES (Fig 1). Cells were incubated under Phe-deprived conditions for 24 h before BPA administration for 2 h. The ^10^B concentrations increased in all of the examined cell lines in a dose-dependent manner, with BPA in the medium under standard culture conditions (Fig 1A and S1 Fig). Phe deprivation in the medium significantly increased the ^10^B concentrations of the SAS, U87-MG, and PANC-1 cells relative to those under standard conditions (Fig 1A). The effect of Phe deprivation was pronounced in SAS cells, where BPA uptake increased by > 4-fold. However, A375 and HaCaT cells showed no increase in ^10^B concentrations following Phe deprivation. These results indicate that pretreatment with Phe-deprived medium enhances BPA uptake in specific tumor cell lines.

**Fig. 1.**
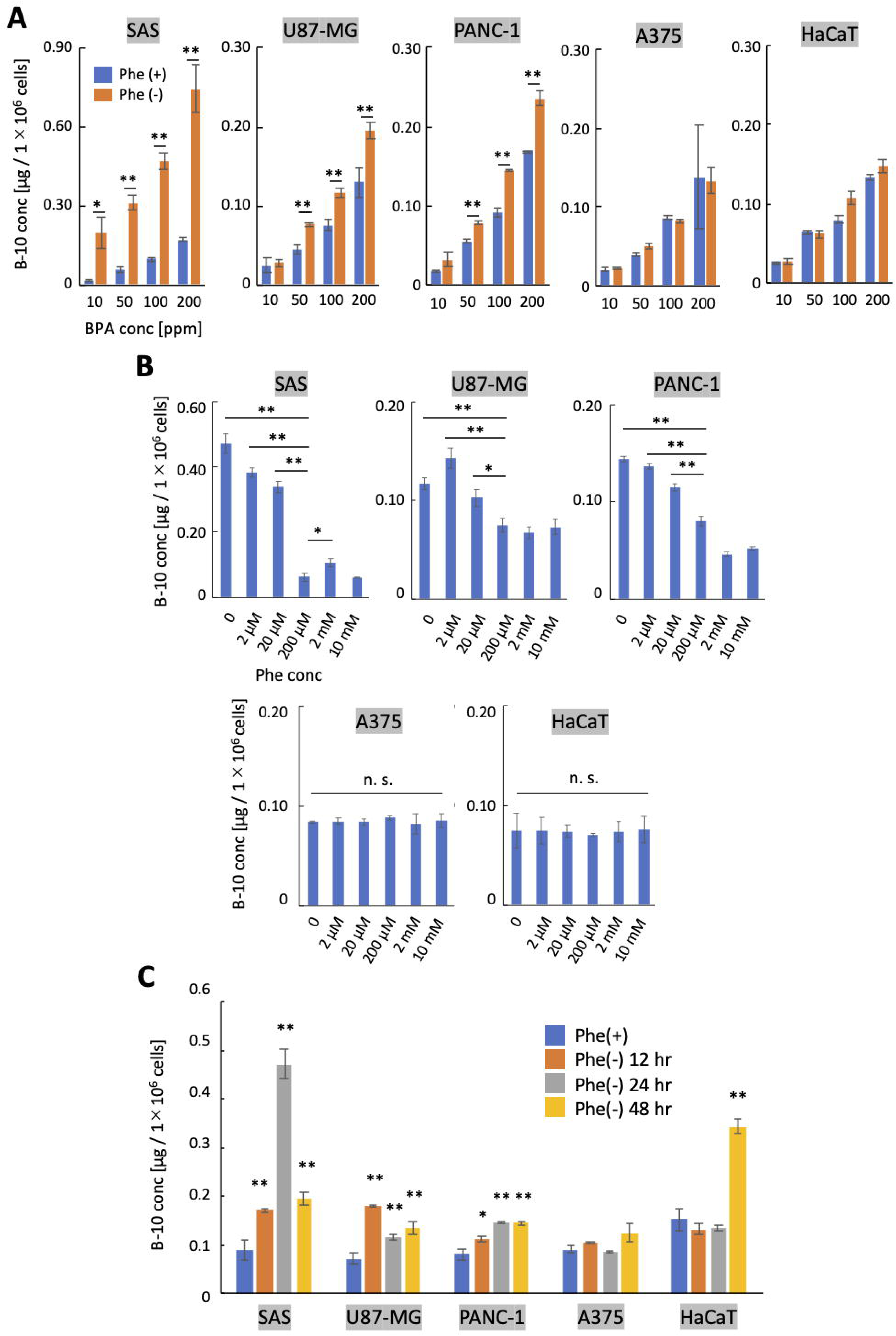
Phenylalanine restriction enhances BPA uptake in tumor cells. (A) B-10 (^10^B) concentrations in the SAS, U87-MG, PANC-1, A375, and HaCaT cells treated with 10–200 ppm BPA. The blue and orange bars represent 24-h pretreatment in normal and Phe-free media, respectively. (B) B-10 concentrations in the cells cultured with 0–10 mM Phe. (C) B-10 concentrations in the control (blue) and cells in Phe-free medium for 12 h (orange), 24 h (gray), and 48 h (yellow). *p < 0.01, **p < 0.001 (Student’s t-test).

To determine whether BPA uptake varies with the amount of Phe in the culture medium, each cell line was cultured for 24 h under different Phe concentrations (Fig 1B). SAS, U87-MG, and PANC-1 cells exhibited higher ^10^B accumulation at 0–20 µM Phe conditions than those at 200 µM–10 mM. The highest ^10^B accumulation was observed under Phe-free conditions in SAS and PANC-1 cells, and at 2 µM Phe in U87-MG cells. SAS cells were highly responsive to low Phe concentrations, whereas A375 and HaCaT cells did not respond to changes in Phe concentration. These findings reveal the differences in responsiveness to a Phe diet among cell lines.

To examine time-dependent changes in BPA uptake under Phe-deprived conditions, concentrations were measured at different incubation times (Fig 1C). In SAS and PANC-1 cells, ^10^B levels peaked after 24 h, whereas in U87-MG cells, the maximum was reached at 12 h. No significant changes were detected in A375 cells at any time point. In HaCaT cells, BPA uptake became evident after 48 h. These observations indicate that time-dependent changes in BPA uptake under Phe deprivation differ among cell lines.

### Cell proliferation of tumor cell lines under Phe-restricted conditions

To evaluate the effects of a low-Phe medium on the proliferation of SAS, U87-MG, PANC-1, A375, and HaCaT cells, cell numbers were counted over 48 h of culture under Phe-restricted conditions (S2A Fig). SAS, A375, and HaCaT cells showed significantly reduced proliferation after 24 h of Phe deprivation (S2B, E, F Figs), whereas U87-MG and PANC-1 cells showed no significant changes (S2C, D Figs). The proliferative responses of these cell lines to Phe restriction varied over 48 h.

### Phe deprivation enhances the efficacy of BNCT on tumor cells

We confirmed the effect of Phe restriction on BNCT by assessing the survival of SAS, U87MG, and PANC-1 cells using the colony formation assay after neutron irradiation in a research reactor (Figs 2A and B). The survival fraction of the BPA-treated plus neutron-irradiated group (Phe(+)_BPA 100ppm) was significantly lower than that of the neutron-irradiated only group (neutron only) (Fig 2C). The survival fractions of the groups with Phe diet and BPA treatment (Phe(–)_BPA 50 ppm and Phe(–)_BPA 100 ppm, respectively) were lower than those of the group without Phe diet (Phe(+)_BPA 100 ppm). The Phe(–)_100 ppm group showed greater reduction across all cell lines, especially SAS, and this was correlated with ^10^B accumulation (Fig 1). These results demonstrate that Phe restriction enhanced BNCT efficacy in the tested cell lines.

**Fig. 2.**
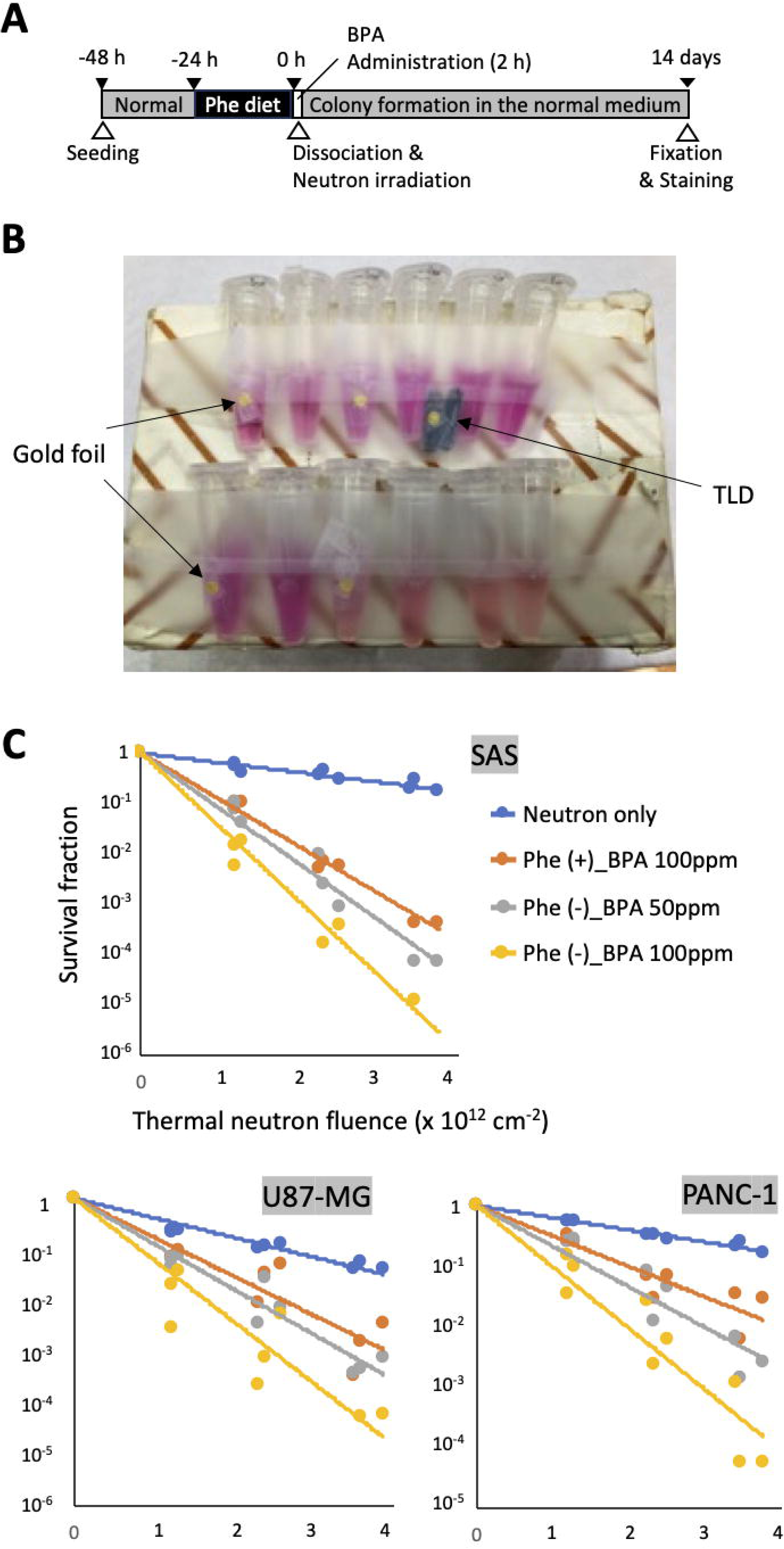
Phenylalanine deprivation enhances the efficacy of neutron irradiation. (A) Schematic representation of Phe restriction, BPA administration, neutron irradiation, and colony formation assay. (B) Sample setup during irradiation. Gold foil and TLD were used to measure the neutron fluence and gamma dose in the 1.5-mL tubes containing the cell suspensions. (C) Survival fractions of the SAS, U87MG, and PANC-1 cell lines. Blue: Neutron only. Orange: 100 ppm BPA. Gray/Yellow: 50/100 ppm BPA with Phe deprivation.

### Phe deprivation induces the integrated stress response and nuclear localization of ATF4

The transcriptomes of SAS, U87-MG, PANC-1, A375, and HaCaT cells were analyzed via RNA sequencing (RNA-seq) after 24 h of Phe deprivation to investigate the side effects of Phe-restricted pretreatment. Genes with significant two-fold changes in TPM values were extracted and compared across the four cell lines. The results indicated that many of the alterations were unique to the individual cell lines, reflecting their different cellular characteristics. However, among all cells, 44 and 52 gene sets were commonly upregulated and downregulated, respectively (Fig 3A and S3A Fig; Gene Lists 1 and 2).

**Fig. 3.**
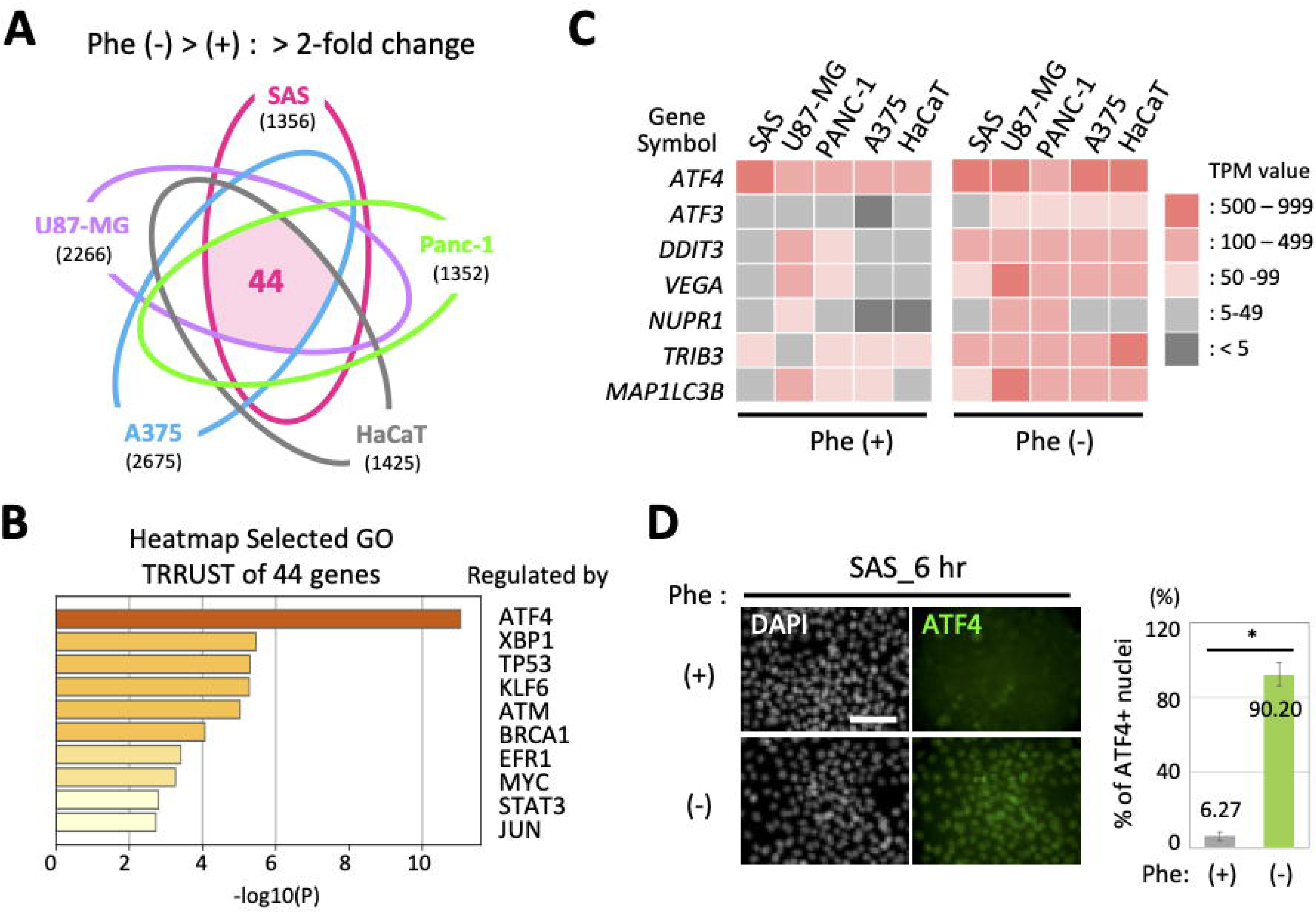
ATF4 stress response to phenylalanine deprivation. (A) Forty-four genes were downregulated in the SAS, U87-MG, PANC-1, A375, and HaCaT cells after 24 h of Phe deprivation. (B) GO TRRUST analysis using Metascape revealed enrichment of ATF4 target genes. (C) Heatmap of the TPM values for ATF4 and its six target genes under control Phe(+) and deprived Phe(‒) conditions. (D) Left panels: ATF4 immunofluorescence in SAS cells under control Phe(+) and Phe-deprived (‒) conditions (6 h). Bar = 100 µm. Right bar graph: Percentage of ATF4-positive nuclei in SAS cells after 6 h of Phe(+) and Phe(‒) conditions (n = 4). * p < 0.0001 (Student’s t-test).

GO analysis of the 44 upregulated genes using Metascape (22) revealed significant enrichment of six direct target genes of the transcription factor ATF4, which regulates the transcription of specific genes during the integrated stress response (ISR) (Fig 3B, C; Gene List 1). *ATF4* mRNA is transcribed in response to cellular stresses, and ATF4 proteins accumulate in the nucleus to bind as heterodimers to the cis-elements of direct target genes (23–25). ATF4 induction has been reported following serine and glycine withdrawal from tumor cell lines (10). We confirmed ATF4 accumulation in the nuclei under Phe deprivation using immunofluorescence (Fig 3D and S3C, D Figs) and calculated the percentage of ATF4-positive nuclei at 6 h after the change to fresh medium. The percentages of cells with ATF4-positive nuclei in the Phe(+) medium were less than 10% for SAS, U87-MG, and HaCaT cells and more than 30% for PANC-1 and A375 cells. However, all of the tested cell lines had a positivity rate of approximately 90% under the Phe(–) condition. These results indicate that Phe deprivation induces ISR in the tested cell lines, with nuclear accumulation of ATF4 apparent from the early stages of treatment.

GO analysis further revealed the enrichment of genes regulated by E2F1 (26) and SP1 (27) among the 52 downregulated genes (S3B Fig and Gene List 2). However, the enrichment scores were low, and E2F1 and SP1 are ubiquitously expressed as housekeeping transcription factors. Thus, we consider that these genes were repressed by ISR at the transcriptional level.

### LAT1 expression levels under Phe deprivation

The RNA-seq data revealed consistent expression of *SLC7A5* coding LAT1 and its coupling factor *SLC3A2* coding 4F2HC across the transporter genes for Phe in all cell lines (S4A Fig). No fluctuations in the expression of other transporter genes were observed under Phe(–) conditions, except for *SLC6A15* in A375 cells. These findings indicate that LAT1/4F2HC is the predominant transport pathway for these cell lines. *SLC7A5* mRNA levels increased in all cell lines except SAS cells in response to Phe deprivation (S4A, B Figs). However, the normalized LAT1 protein expression levels were largely unchanged under the Phe(+) and Phe(–) conditions (S4C Fig). The transcriptional alteration of *SLC7A5* did not directly affect LAT1 levels in these cells within the framework of the ISR to Phe deprivation.

### Phe deprivation alters the intracellular amino acid levels of tumor cell lines

The intracellular amino acid levels were validated in SAS, U87-MG, and A375 cells, which showed different proliferation and BPA uptake responses to Phe deprivation. The amino acid levels of the cells under the Phe(–) and Phe(+) conditions were normalized based on the results of LC-MS/MS analysis of water-soluble primary metabolites (Fig 4A). The cellular Phe levels of SAS and A375 cells decreased under deprivation, but those of U87-MG cells were unchanged, suggesting the presence of mechanisms to buffer Phe. Increased levels of several amino acids, including substrates for the LAT1/4F2HC complex, were observed in the SAS and U87-MG cells under Phe(–). However, the level of aspartic acid decreased in the U87-MG cells. The amino acid levels of A375 cells remained stable under Phe deprivation, except for those of cysteine, glutamic acid, and proline, which decreased, and of asparagine, which increased. These findings suggest that deprivation of a particular amino acid does not simply induce its shortage or increase the uptake of other amino acids, but that the response pattern varies depending on the cell line.

**Fig. 4.**
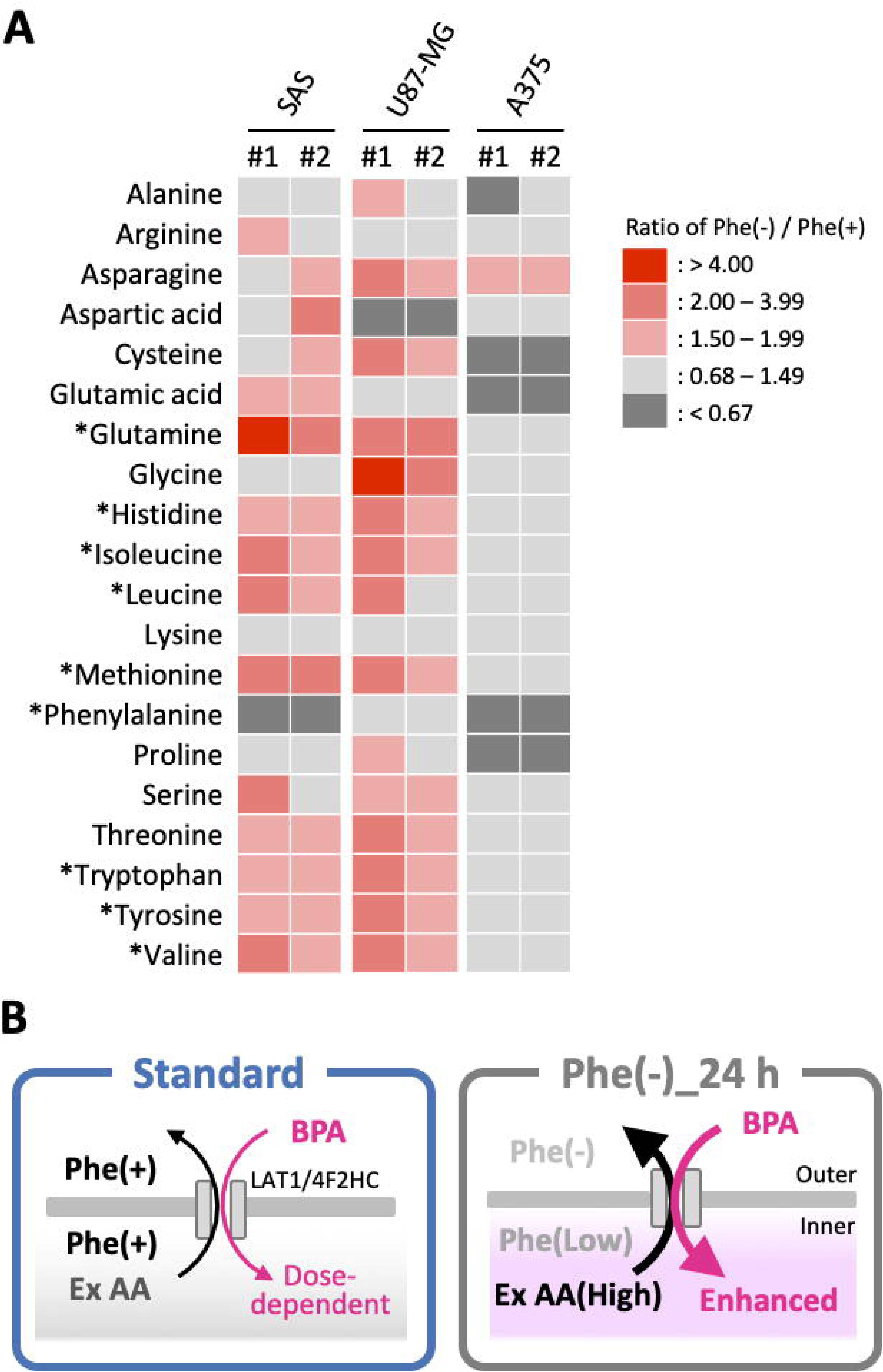
Phenylalanine deprivation alters cellular amino acid levels. (A) Heatmap showing ratios of amino acid levels in SAS, U87-MG, and A375 cells under Phe(‒) and Phe(+) conditions for 24 h (#1 and #2: replicates). *Substrates of the LAT1/4F2HC amino-acid transporter (AAT). (B) BPA uptake model via AAT under standard Phe(+) (left) and Phe(‒) (right) conditions. Ex AA: Exchange amino acids.

## Discussion

We demonstrated *in vitro* that Phe deprivation enhanced BPA uptake and improved BNCT efficacy in specific tumor cell lines, highlighting the potential of short-term nutrient restriction as a preconditioning strategy to enhance treatment effectiveness.

Our working model suggests that BPA uptake under Phe restriction is regulated by the balance between intracellular Phe and other amino acids that can act as exchange substrates for the LAT1/4F2HC complex (Fig 4B and S5 Fig). Under standard culture conditions, the cells contain sufficient amino acids and take up BPA from the medium in a dose-dependent manner (S1 Fig). SAS and U87-MG cells likely increase amino-acid uptake via LAT1/4F2HC under Phe restriction, resulting in elevated intracellular levels of exchangeable substrates that could promote BPA transport. The intracellular Phe levels decreased, which enhanced the uptake of BPA, a Phe analog. The mechanism remains unclear, but U87-MG cells maintained their intracellular Phe levels, which limited the increase in BPA uptake to a moderate extent. In contrast, A375 cells exhibited a reduction in their intracellular Phe levels without a compensatory increase in alternative exchange substrates, and their BPA uptake remained dose-dependent, as observed under standard conditions.

Previous studies have reported the efficacy of amino acid pretreatment in enhancing the T/N ratio of BPA. Watanabe *et al*. reported that preloading with Phe at high concentrations reduced BPA accumulation in the normal tissues of tumor-bearing mice and increased the T/N ratio (12). Preloading with tyrosine and L-DOPA, both LAT1 substrates, has been shown to increase BPA uptake in tumor cells (28–31). Optimizing the T/N ratio for BPA will require systematic evaluation of Phe restriction and LAT1-substrate preloading schedules under controlled dietary conditions.

The potential of amino acid restriction to suppress tumor cell proliferation and improve prognosis has been reported (8‒10). We demonstrated that Phe restriction reduced cell proliferation and induced ISR with nuclear ATF4 accumulation in all of the tested cell lines. However, BPA uptake increased in some cell lines but not in others under Phe restriction, suggesting that the ATF4 response occurs via a distinct pathway from BPA uptake regulation. Our RNA-seq data revealed substantial differences among the cell lines under both standard and Phe-restricted conditions, necessitating further investigations to identify appropriate tumor types for BNCT treatment in combination with a Phe-restricted diet.

Due to the intrinsic heterogeneity of tumor cell lines, their responsiveness to BPA accumulation under Phe-restricted conditions was not uniform. At present, specific tumor types or molecular markers that predict the effectiveness of this pretreatment have not yet been identified. Further investigations are required to elucidate the molecular mechanisms underlying this heterogeneous responsiveness. In addition, establishing a Phe-restricted culture system with commercially available normal human cells was difficult due to limited sample availability and undisclosed medium compositions. Consequently, their responsiveness to Phe restriction could not be evaluated in this study. Furthermore, as this study was conducted *in vitro,* the physiological effects of Phe restriction should be thoroughly validated *in vivo* in future experiments.

This study supports that Phe restriction can enhance the tumor-killing effects of BNCT. All of the neutron-irradiated cell lines with increased BPA uptake due to Phe restriction demonstrated a direct correlation between BPA uptake and BNCT outcomes. We propose that PKU dietary therapy may serve as a clinical application for Phe-restricted BNCT. PKU, a congenital deficiency of phenylalanine hydroxylase, leads to delayed mental development and epilepsy, and dietary restriction is the standard treatment. Short-term Phe restriction has minimal side effects and may be implemented as pretreatment for BNCT. Further validation of these *in vitro* findings using animal models will enable us to advance this approach toward clinical adoption.

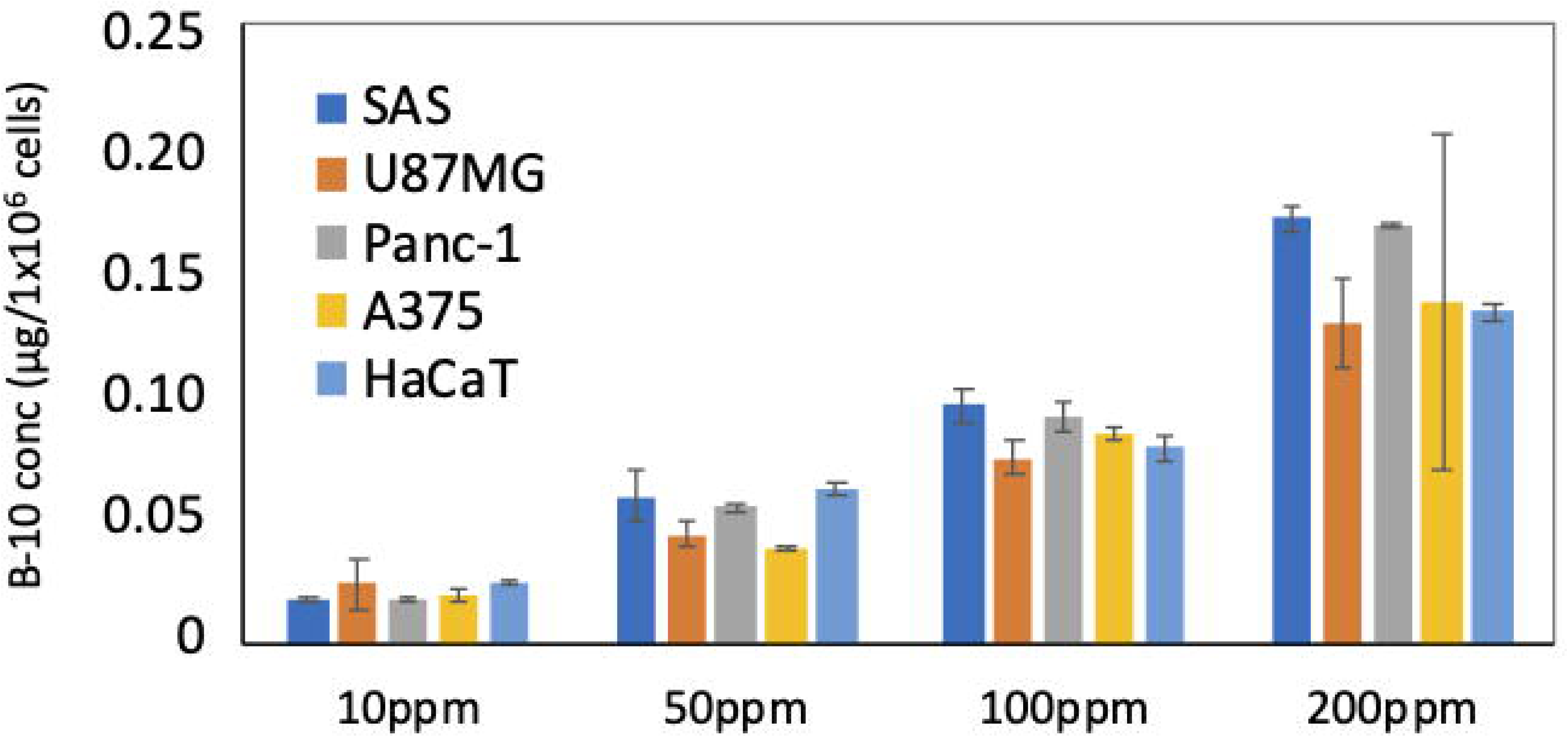

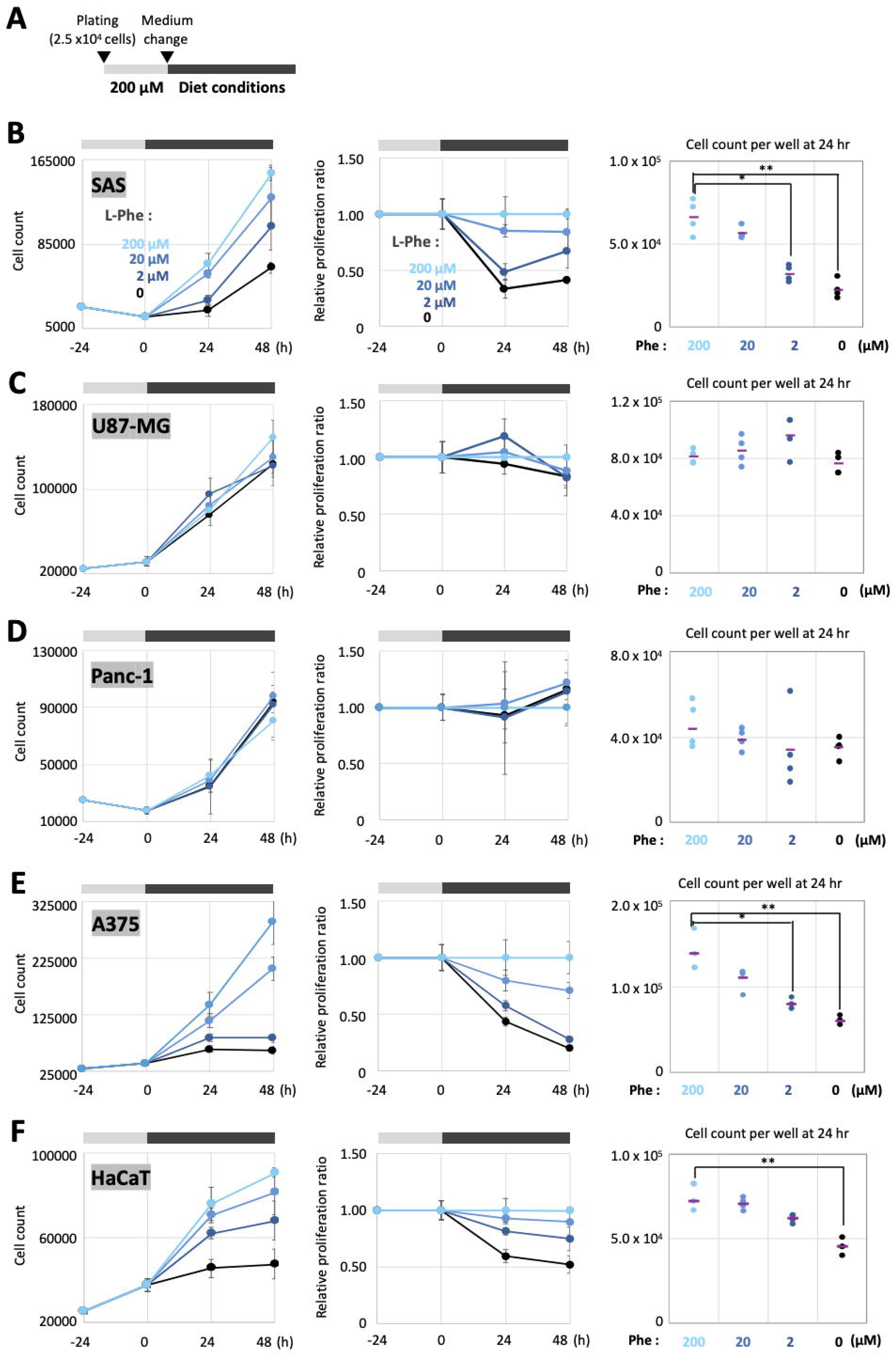

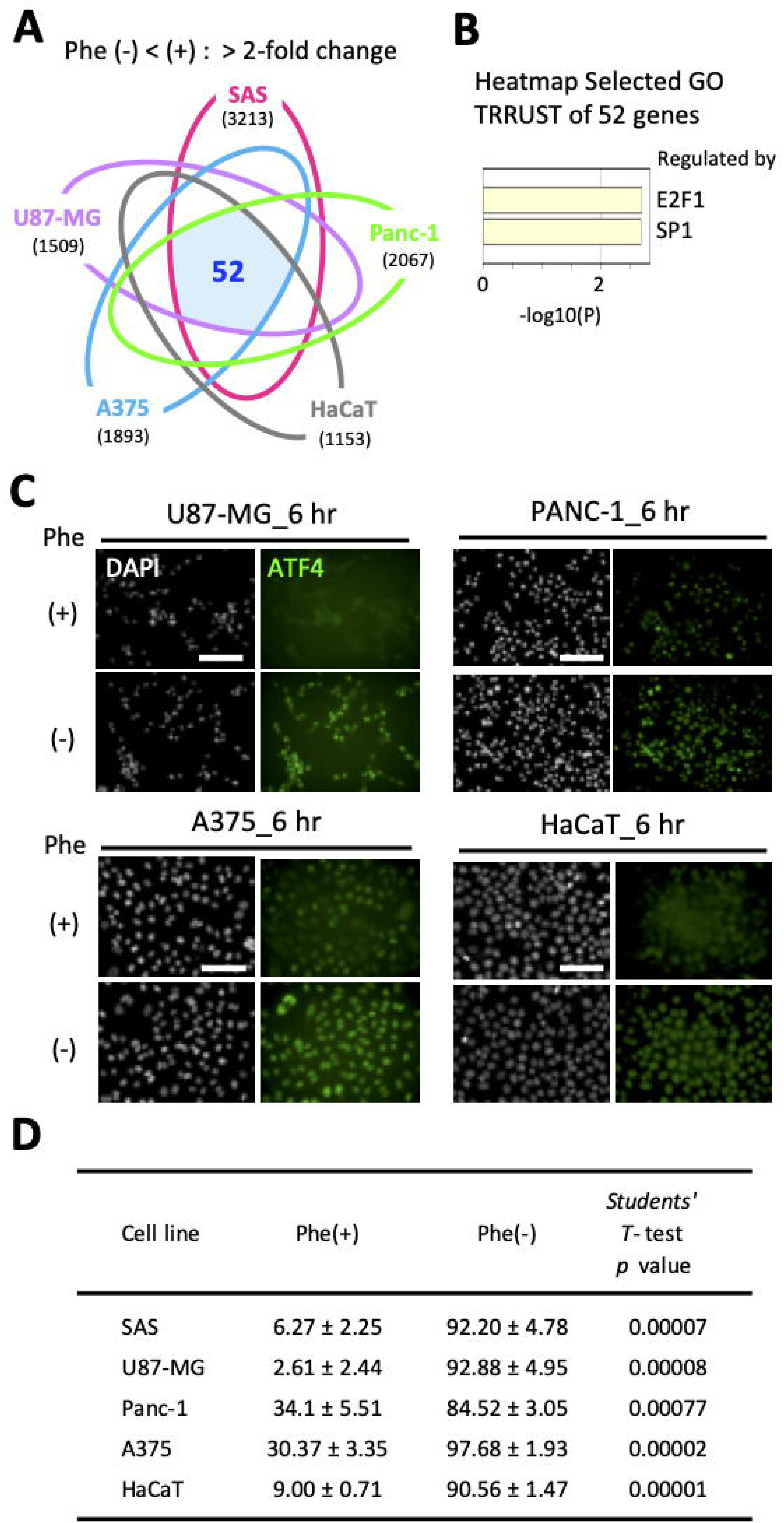

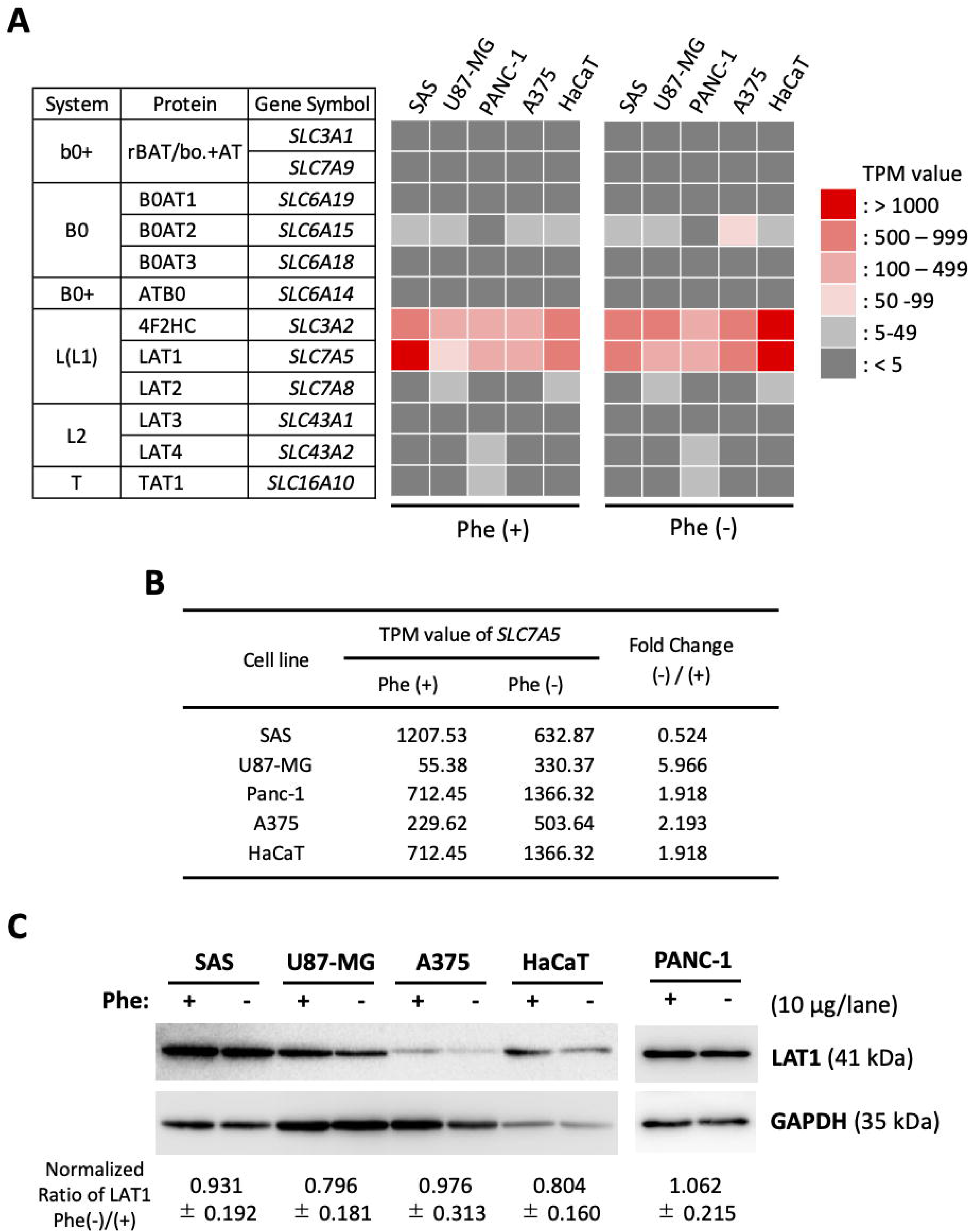

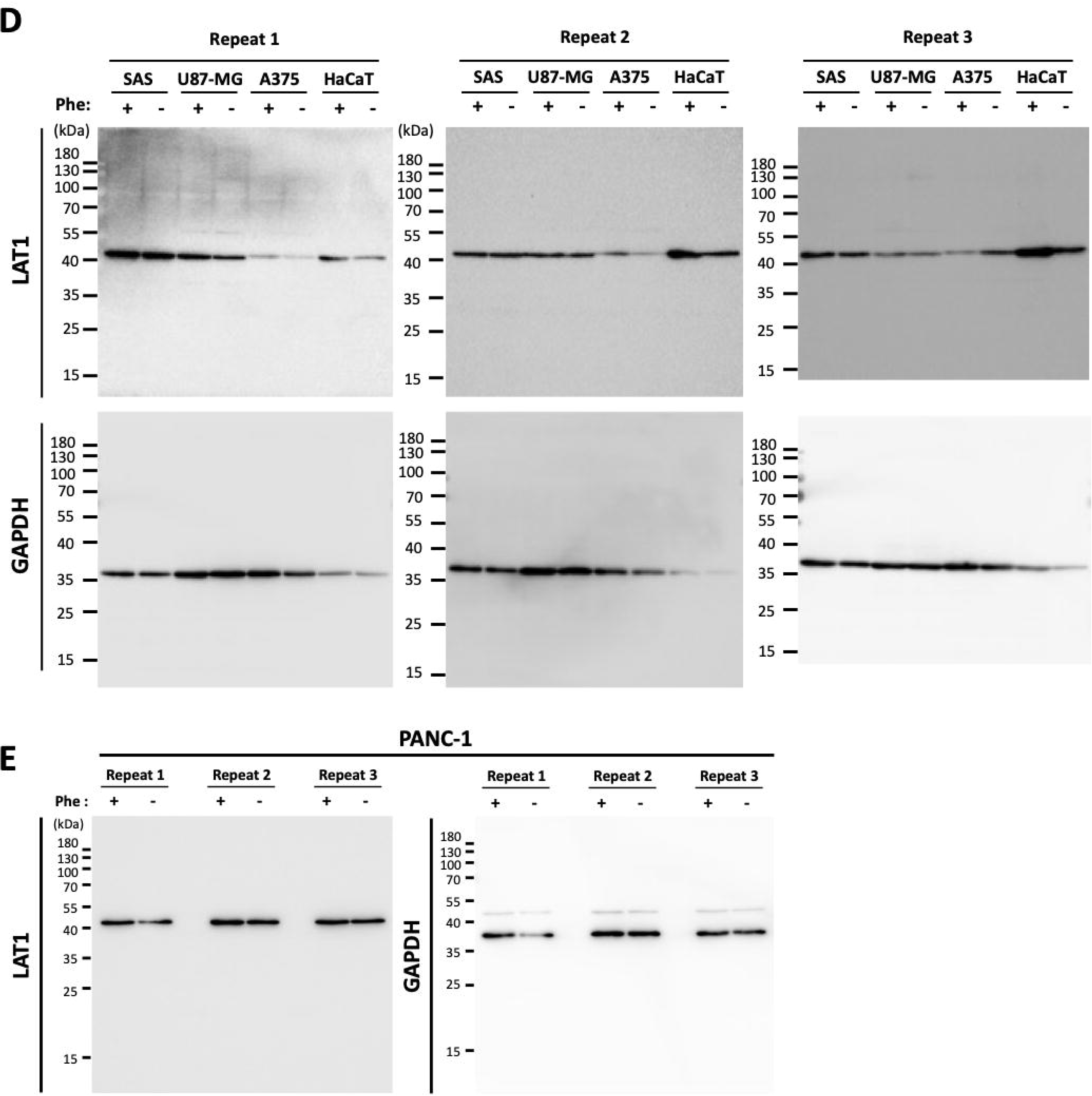

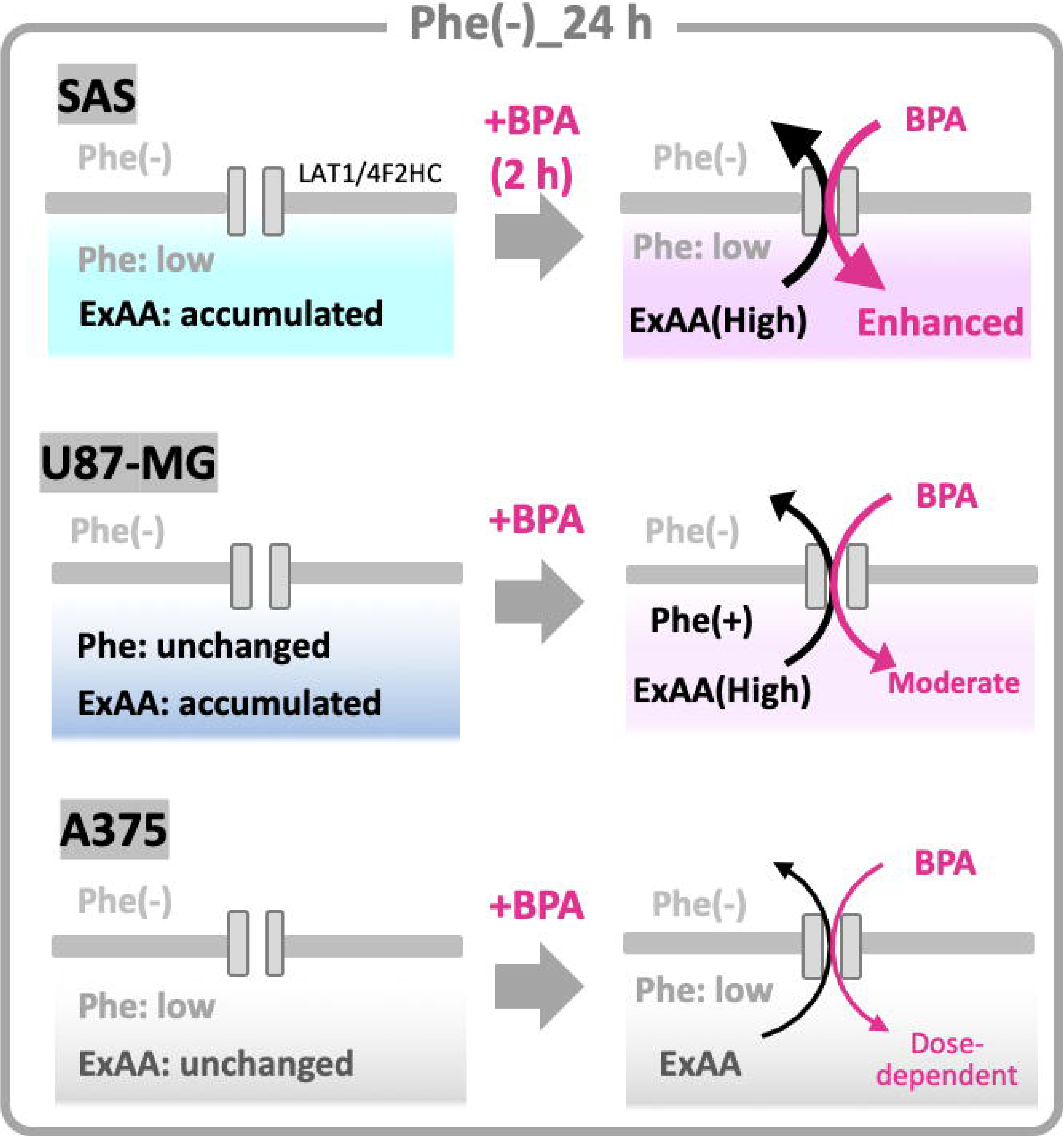

## Supporting information

Supplementary Procedures

Gene List 1

Gene List 2

## Acknowledgments

We thank Norito Kato and Hiroshi Takenaka (KPUM, Japan), the NGS Core Facility (RIMD, Osaka University, Japan), Mari Adachi and Takeshi Okumura (Shimadzu Techno-Research Inc., Japan), and SiC Engineering Inc./J-BEAM (Fukushima, Japan) for their technical support and helpful discussions. The authors used DeepL Write (DeepL SE) and ChatGPT (OpenAI) to assist in editing the language of the manuscript, and all content was reviewed and verified by the authors.

