## Supplementary Procedures for "L-Phenylalanine restriction amplifies boron neutron capture therapy efficacy through increased L-boronophenylalanine uptake and induces activating transcription factor 4 stress response in tumor cell lines"

**Supplementary procedure**

**Cell counting for the Phe diet**

The cells ($2.5 \times 10^4$) were plated in 12-well plates containing control medium. The medium was replaced with diet conditions containing 0–10 mM Phe (13039-62, Nacalai Tesque) 24 h later. The cells were dissociated using 0.25% trypsin-ethylenediaminetetraacetic acid (EDTA) solution (T2605, Sigma-Aldrich, St. Louis, MO, USA), suspended in the medium, and counted per well using Trypan Blue Solution (207-17081, FUJIFILM Wako) and a Cell Counter Plate (177-112C, Watson Co. Ltd., Tokyo, Japan). The mean values for the four wells were plotted.

**Western blotting**

The cells were dissociated using trypsin-EDTA, washed with PBS, and lysed using RIPA buffer (150 mM NaCl, 0.1% Triton X-100, 0.5% sodium deoxycholate, 0.1% sodium dodecyl sulphate, and 50 mM Tris-HCl [pH 8.0]), Protease Inhibitor Cocktail (03969-21, Nacalai Tesque), and Phosphatase Inhibitor Cocktail (07575-51, Nacalai Tesque). The protein content of the lysate was determined using the CBB protein assay solution (29449-44, Nacalai Tesque). Ten micrograms of the lysate were boiled with Laemmli sample buffer (2% SDS, 5% 2-mercaptoethanol, 10% glycerol, 0.002% bromophenol blue, and 0.0625 M Tris-HCl [pH 6.8]) and resolved by SDS-PAGE. The proteins were transferred to a PVDF membrane (10600058, Amercham, Buckinghamshire, UK). These were visualized by immunoblotting using mouse anti-LAT1 (1:1000; sc-374232, Santa Cruz Biotechnology, Dallas, TX, USA) and mouse anti-GAPDH (1:1000; 5A12, FUJIFILM Wako) primary antibodies, HRP-conjugated horse anti-mouse IgG (1:5000; 7076, Cell Signaling Technologies, Danvers, MA, USA) secondary antibody, Chemi-Lumi One Super (02230-14, Nacalai Tesque), and ChemiDoc Touch MP (17001402JA, Bio-Rad, Hercules, CA, USA). Signal intensities for LAT1 and GAPDH were determined using ImageJ (https://imagej.net/ij/).
