## Supplementary Figure Captions for "L-Phenylalanine restriction amplifies boron neutron capture therapy efficacy through increased L-boronophenylalanine uptake and induces activating transcription factor 4 stress response in tumor cell lines"

**Fig. S1 Dose-dependent BPA uptake by tumor cell lines under standard culture conditions.** BPA uptake by SAS, U87MG, PANC-1, A375, and HaCaT cells cultured in the control medium. The cell lines were administered BPA at concentrations of 10, 50, 100, and 200 ppm and ashed after 2 h of incubation.

**Fig. S2 Reduction of tumor cell proliferation with a low Phe diet.**

(A) Time course of cell culture on the Phe diet (0, 2, 20, and 200 μM). (B–F) Graphs showing the cell count per well from −24 to 48 h at 24-h intervals (left), relative proliferation ratio compared with the control (200 μM Phe) (middle), and cell count per well at 24 h for SAS (B), U87-MG (C), PANC-1 (D), A375 (E), and HaCaT (F) cells. * and ** indicate $p < 0.01$ and $p < 0.001$, respectively, based on Student's t-test.

**Fig. S3 Gene subtraction and ATF4 response following Phe deprivation in tumor cell lines.**

(A) Fifty-two genes were downregulated in SAS, U87-MG, PANC-1, A375, and HaCaT cells after Phe deprivation for 24 h. (B) GO TRRUST analysis of the 52 genes using Metascape revealed no significant enrichment of the downstream target genes. (C) Immunofluorescence for ATF4 in U87-MG, PANC-1, A375, and HaCaT cells after 6 h of normal Phe (+) (upper panels) and Phe deprivation (-) (lower panels). Bar = 100 μm. (D) Average percentages of ATF4-positive nuclei relative to DAPI-positive nuclei in the cell lines after 6 h of Phe(+) and Phe(-) conditions. N = 4.

**Fig. S4 LAT1 expression under Phe deprivation.** (A) Heatmap showing the TPM values of the SLC genes related to Phe uptake based on the RNA-seq data for SAS, U87-MG, PANC-1, A375, and HaCaT cell lines. The data for the normal Phe(+) and Phe(-) diets are provided in the left and right panels, respectively. (B) Summary of the TPM values of the *SLC7A5* for each cell line. (C) Western blotting results for LAT1 (41 kDa) and GAPDH (35 kDa) for each cell line under Phe(+) and Phe(-) conditions for 24 h. The ratios of the LAT1 expression levels normalized to those of GAPDH (Phe(-) to Phe(+)) are provided (N = 3). (D, E) Western blotting results for LAT1 and GAPDH for the SAS, U87-MG, A375, and HaCaT cell lines (D) and PANC-1 cell line (E).

**Fig. S5 Exchange substrates of LAT1 and BPA following Phe deprivation in tumor cell lines.** Schematic models of the regulation of BPA uptake in SAS, U87-MG, and A375 cells after Phe deprivation for 24 h. The balance of intracellular Phe and exchange amino acid substrate (ExAA) concentrations may serve as a regulatory factor for the increased BPA uptake observed following Phe deprivation.
